# Impaired Lef1 activation accelerates iPSC-derived keratinocytes differentiation in Hutchinson-Gilford Progeria Syndrome

**DOI:** 10.1101/2022.02.22.481406

**Authors:** Xiaojing Mao, Zheng-Mei Xiong, Huijing Xue, Markus A. Brown, Yantenew G. Gete, Reynold Yu, Linlin Sun, Kan Cao

**Affiliations:** Department of Cell Biology and Molecular Genetics, University of Maryland, College Park, MD 20817, USA; National Human Genome Research Institute, National Institutes of Health, Bethesda, MD, USA; Key Laboratory of Diagnosis and Treatment of Aging and Physical-chemical Injury Diseases of Zhejiang Province, The First Affiliated Hospital, School of Medicine, Zhejiang University, Hangzhou, Zhejiang, China

## Abstract

Hutchinson-Gilford progeria syndrome (HGPS) is a detrimental premature aging disease caused by a point mutation in the human *LMNA* gene. This mutation results in the abnormal accumulation of a truncated pre-lamin A protein called progerin. Among the drastically accelerated signs of aging in HGPS patients, severe skin phenotypes such as alopecia and sclerotic skins always develop with the disease progression. Here, we study the HGPS molecular mechanisms focusing on early skin development by differentiating patient-derived induced pluripotent stem cells (iPSCs) to a keratinocyte lineage. Interestingly, HGPS iPSCs showed an accelerated commitment to the keratinocyte lineage than the normal control. To study potential signaling pathways that accelerated skin development in HGPS, we investigated the WNT pathway components during HGPS iPSCs-keratinocytes induction. Surprisingly, despite the unaffected β-catenin activity, the expression of a critical WNT transcription factor LEF1 was diminished from an early stage in HGPS iPSCs-keratinocytes differentiation. Chromatin immunoprecipitation (ChIP) experiment further revealed strong bindings of LEF1 to the early-stage epithelial developmental markers K8 and K18 and that the LEF1 silencing by siRNA down-regulates the K8/K18 transcription. During the iPSCs-keratinocytes differentiation, correction of HGPS mutation by Adenine base editing (ABE), while in a partial level, rescued the phenotypes for accelerated keratinocyte lineage-commitment. ABE also reduced the cell death in HGPS iPSCs-derived keratinocytes. These findings brought new insight into the molecular basis and therapeutic application for the skin abnormalities in HGPS.

## Introduction

Hutchinson-Gilford Progeria syndrome (or HGPS, progeria) is an ultra-rare but devastating genetic disorder that causes premature aging in children (Eriksson et al., 2003). The HGPS patients appear normal at birth but soon develop a series of severe health conditions that occur in the process of aging, including alopecia, scleroderma, subcutaneous fat loss, bone and joint defects, cardiovascular diseases, etc. Most HGPS patients die of arteriosclerosis at an average age of 14.5 (Gordon, Brown, & Collins, 2019; Merideth et al., 2008). Classical HGPS is caused by a *de novo* point mutation within the human *LMNA* gene (c.1824C>T). This gain-of-function mutation activates a cryptic splice site which generates a truncated, unproperly processed lamin A precursor that retains a hydrophobic farnesyl tail named progerin (De Sandre-Giovannoli et al., 2003; Eriksson et al., 2003). The accumulation of progerin in the cell nucleus induces abnormal nuclear morphology and altered gene expression and chromatin structure (Goldman et al., 2004; McCord et al., 2013; Shumaker et al., 2006). However, how progerin accelerates the aging process in many tissues and organs during their development remains largely unclear, particularly in the skin, the human body’s largest organ.

Human skin consists of the epidermis, dermis, and subcutaneous tissues (Millington, 1983). In HGPS, skin abnormalities, including reduced epidermis thickness, alopecia, dimpling, wrinkling, altered pigmentation, and decreased subcutaneous fat, overlap significantly with signs of normal aging (Gordon et al., 2019; Merideth et al., 2008). As the outermost surface of human skin, the epidermis is susceptible to environmental stimuli and develops signs of aging. Keratinocytes are the primary cell type in the stratified epidermis. Moving outward from the basement membrane, the basal layer (*Stratum basale*) keratinocytes undergo a sequential differentiation process to form the spinous layer (*Stratum spinosum*), granular layer (*Stratum granulosum*), and cornified layer (*Stratum corneum*) above (Fuchs & Raghavan, 2002).

HGPS skin abnormalities have been studied in transgenic mice with inducible epidermal expression of progerin under the control of Keratin 5 (K5) or Keratin 14 (K14) promotors (Sagelius et al., 2008; Wang et al., 2008). Postnatal progerin expression under K5 resulted in premature exhaustion of adult stem cells in mouse skins (Rosengardten, Mckenna, Grochová, & Eriksson, 2011). Epidermal keratinocytes with aberrant proliferation capacity and altered gene expression have also been found in these animals (Rosengardten et al., 2011; Sagelius et al., 2008). In the same animal model with embryonic induction of progerin expression, keratinocytes from the interfollicular epidermis (IFE) presented an abnormal tendency to divide symmetrically, generating excessive basal keratinocytes at the basement membrane. As a result, basal cells were often spotted in suprabasal layers of the epidermis in HGPS transgenic mice and were likely to induce epidermal hyperplasia (Mckenna et al., 2014; Sagelius et al., 2008; Sola-Carvajal et al., 2019). Pathway analysis indicated that impaired WNT/β-catenin signaling might be responsible for this phenomenon, as progerin disrupted the nuclear localization of emerin and nesperin-2, impeding the nuclear transportation of β-catenin (Choi et al., 2018; Sola-Carvajal et al., 2019). While these transgenic animals with lineage-specific progerin expression served as great models to study skin abnormalities in HGPS, they may not fully represent the symptoms in HGPS patients due to genetic divergence between humans and mice. Moreover, these studies focused on keratinocytes from fully developed mouse skins, and thus only those phenotypes at late developmental stages were characterized. To investigate the HGPS disease progression in early epidermal development with a humanized model, we took advantage of the patient-derived iPSCs and monitored the full differentiation process towards a keratinocyte lineage.

Currently, there is no cure for HGPS. Yet, the recent application of Adenine Base Editing (ABE) in HGPS primary cells and an animal model offered up-and-coming results (Koblan et al., 2021). This novel gene-editing tool linked a deoxyadenosine deaminase to a deactivated CRISPR vector, which conducts A•T to G•C base pairs conversion at the targeted genome region (Gaudelli et al., 2017). In HGPS, with the precise-targeting single-guide RNA (sgRNA), these base editors directly corrected the *LMNA* c.1824C>T mutation with high efficiency. *In vitro* base editing in HGPS primary fibroblasts and endothelial cells showed complete progerin clearance, while the embryonic ABE-AAV9 injection greatly improved the health of HGPS mice and significantly extended their lifespans (Gete et al., 2021; Koblan et al., 2021).

In this study, we investigated the early development of keratinocytes in HGPS. We first differentiated a pair of well-characterized normal and HGPS patient-derived induced pluripotent stem cells (iPSCs) into a keratinocyte lineage (Atchison et al., 2020; Atchison, Zhang, Cao, & Truskey, 2017; Gete et al., 2021; Xiong, LaDana, Wu, & Cao, 2013; H. Zhang, Xiong, & Cao, 2014). We found that HGPS iPSCs showed an accelerated commitment to the keratinocyte lineage during the differentiation process. Furthermore, we identified a possible upstream effector in epidermal development: LEF1, whose activation was diminished in HGPS iPSCs differentiation. By utilizing the novel gene-editing technique ABE, we partially corrected the genetic mutation in HGPS during the keratinocytes induction and rescued the defects in HGPS iPSCs-derived keratinocytes.

## Results

### Differentiating normal and HGPS-derived iPSCs into keratinocytes

To differentiate normal and HGPS patient-derived iPSCs into the keratinocyte lineage *in vitro*, we adapted a four-week differentiation protocol including two inductions with retinoid acid (RA) and BMP-4 at the initial stage, followed by keratinocyte growth medium changes every other day for up to 28 days (Itoh, Kiuru, Cairo, & Christiano, 2011; Kogut, Roop, & Bilousova, 2014). At the end of differentiation, a subpopulation of basal keratinocyte-like cells with cobblestone morphology was observed (Fig 1A). To evaluate the efficacy of differentiation, we first detected the expression of lamin A/C, progerin, and basal keratinocyte markers ΔNp63 and K14 by immunofluorescence staining (Fig 1B-C). As expected, all these iPSCs-derived cells showed ΔNp63 and K14 expression with different fluorescence intensities. Their cellular distribution was comparable to those in primary keratinocytes (Fig 1C). Progerin expression was only detected in cells differentiated from HGPS iPSCs (Fig 1B).

**Figure 1.**
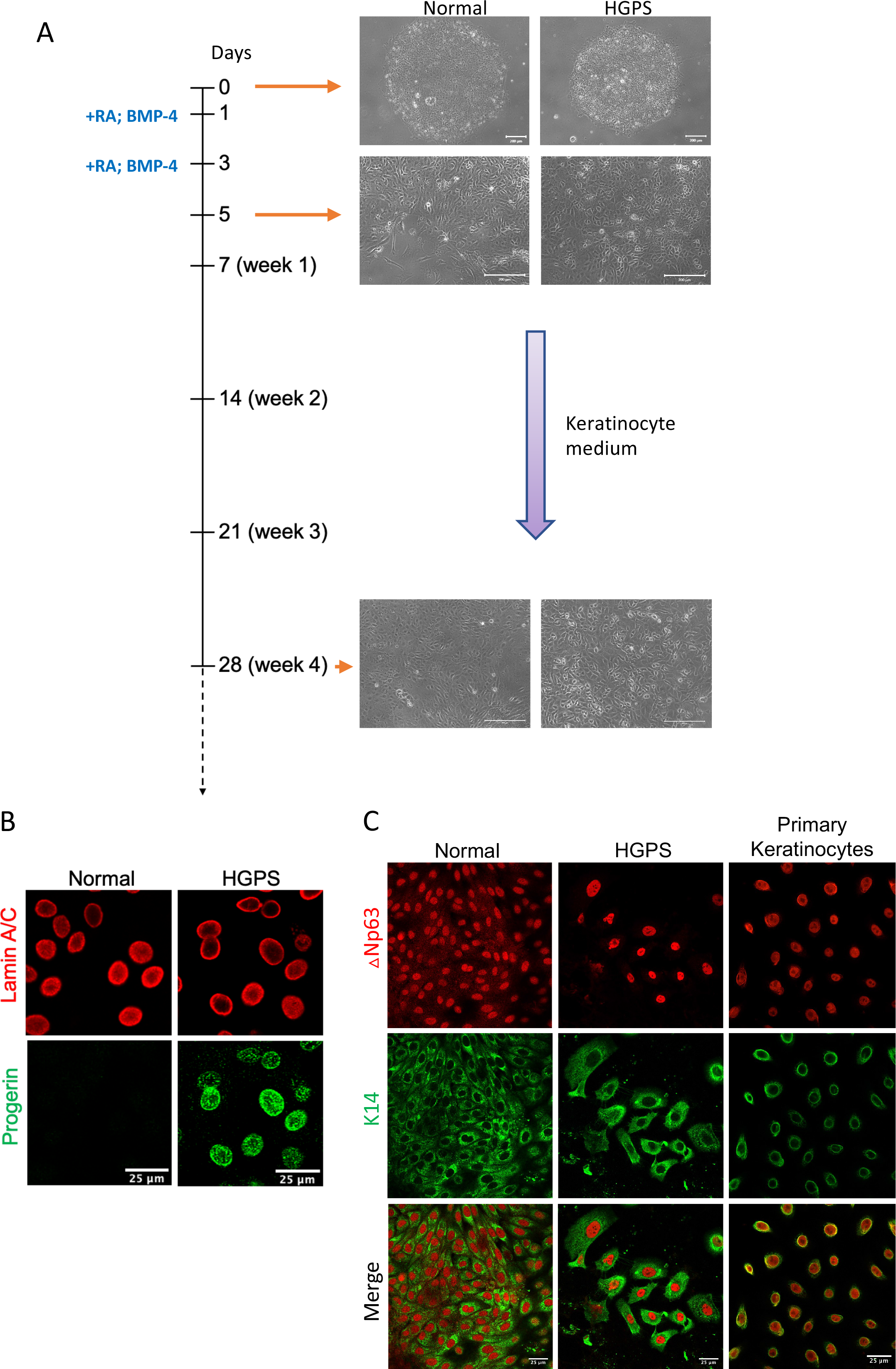
Differentiating normal and HGPS-derived iPSCs into keratinocytes. A. Schematic representation of the protocol for differentiating normal and HGPS patient-derived iPSCs into a keratinocyte lineage. RA and BMP-4 were added on day 1 and day 3. Cells were maintained in keratinocyte growth medium for more than 28 days (4 weeks). Representative phase-contrast images of normal and HGPS iPSCs at different time points during keratinocyte induction were shown next to the timeline (scale bar = 200μm). B. Lamin A/C and progerin expression in normal and HGPS iPSCs differentiated cells at week 4 indicated by immunofluorescence staining (scale bar = 25μm). C. Basal layer keratinocyte markers ΔNp63 and K14 expression in normal and HGPS iPSCs differentiated cells at week 4 as well as in primary keratinocytes indicated by immunofluorescence staining (scale bar = 25μm).

Conventionally, a succession of keratin genes expressed in keratinocytes has been identified during mammalian epidermal development: in the early embryonic stage, the expression of keratin pair K8-K18 peaks in single-layered epithelial progenitors that give rise to epidermis and declines rapidly towards the keratinocyte commitment (Jackson, Grund, Winter, Franke, & Illmensee, 1981). Afterward, with the induction of p63, these progenitor cells are committed to the epidermal fate by turning on the K5-K14 expression (Nagarajan, Romano, & Sinha, 2008; Romano et al., 2012). These p63+, K5/K14+ cells are adult stem cells that can proliferate and differentiate into spinous cells, granular cells, and squames by upward stratification (Fuchs, 2007). One keratin type that marks this terminal differentiation is K1 (Fuchs & Green, 1980; L. Zhang, 2018). To characterize the normal and HGPS iPSCs-keratinocytes differentiation at each differentiation stage, the expression of these markers was quantified weekly for four weeks (Fig 2A-B) with quantitative RT-PCR and Western blotting analysis. We found that Lamin A expression was gradually upregulated during the four-week differentiation and only detected progerin expression in HGPS samples (Fig 2A). The early marker K8 was first detected upon the keratinocytes induction, and interestingly, its expression declined more rapidly in HGPS iPSCs-keratinocytes differentiation compared to the normal situation. Accordingly, basal layer keratinocyte markers ΔNp63 and K14 expression increased through normal and HGPS iPSCs differentiation. At each time point detected, ΔNp63 and K14 levels were consistently higher in HGPS than normal control. The terminal differentiation marker K1 was only seen in HGPS iPSCs-derived keratinocytes at week 4 but absent in normal iPSCs differentiation, suggesting that only the 4-week differentiated population in HGPS contained the fully matured keratinocytes to initiate the terminal differentiation (Fig 2A and B).

**Figure 2.**
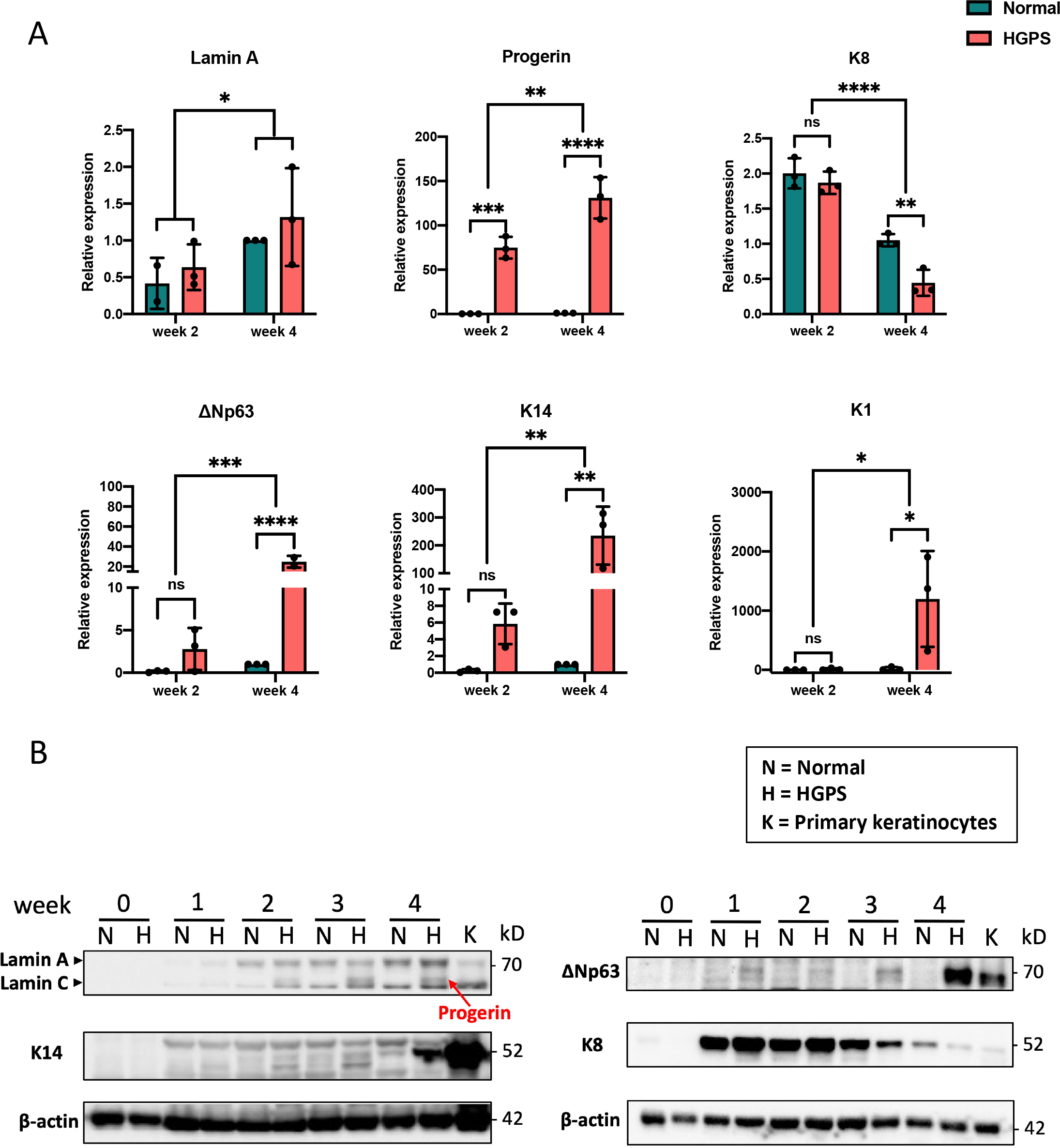
The expression of keratinocyte markers during iPSCs-keratinocytes differentiation. A. Quantitative RT-PCR analysis of the relative expression of Lamin A, progerin, ΔNp63, K14, K8 and K1 during normal and HGPS iPSCs-keratinocytes induction. Data were normalized to endogenous *ACTB* mRNA and to the average of Normal week 4. Data were presented as mean ± SD (n = 3). \**P* < 0.05, \*\**P* < 0.01, \*\*\**P* < 0.001, \*\*\*\**P* < 0.0001, ns, not significant, Two-way ANOVA followed by Tukey’s multiple comparisons test. B. Western blot analysis of Lamin A/C (including progerin in HGPS differentiated cells indicated by the red arrow), ΔNp63, K14, and K8 expression during normal and HGPS iPSCs-keratinocytes induction. Primary keratinocytes were used as a positive control. All experiments were repeated at least three times, and representative data were shown as indicated.

### Dysregulated cell cycle and increased cell death in HGPS iPSCs-keratinocytes differentiation

Progerin disrupts the cell cycle progression in mammalian cells (Cao, Capell, Erdos, Djabali, & Collins, 2007; Dechat et al., 2007; H. Zhang et al., 2014). To investigate the cell cycle partitioning in HGPS epidermal development, we first labeled the normal and HGPS cells at the intermediate (week 2) and late (week 4) iPSCs-keratinocytes differentiation stages using Bromodeoxyuridine (BrdU) as an indication for the DNA synthesis at the S phase. These cells were then stained with fluorescent BrdU antibody and propidium iodide (PI) and quantified for frequencies at each cell cycle stage by a flow cytometry analysis (Fig EV1A and B, Fig 3A). Notably, we observed a more significant fraction of HGPS cells at the G2/M phase than normal differentiation at the intermediate stage. This disparity became more significant at the late stage of differentiation (Fig 3A).

**Figure 3.**
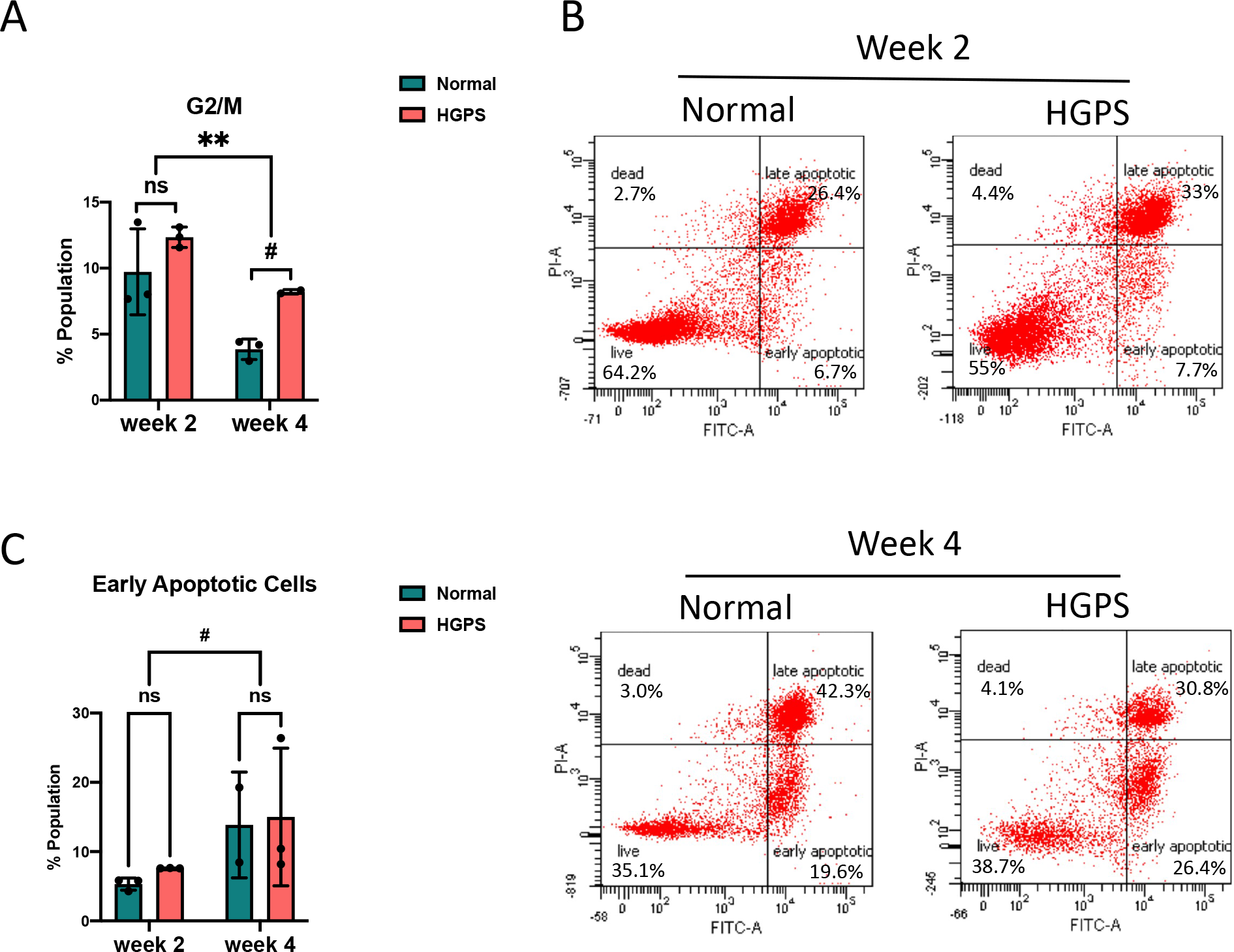
Cell cycle and apoptosis during iPSCs-keratinocytes differentiation. A. Quantification of the G2/M phase cell cycle partitioning during normal and HGPS iPSCs-keratinocytes induction. Data were presented as mean ± SD (n = 3). #P < 0.1, **P < 0.01, ns, not significant, Two-way ANOVA followed by Tukey’s multiple comparisons test. B. Representative flow cytometry plots showing PI-annexin V apoptosis assay during normal and HGPS iPSCs-keratinocytes induction. The gates were set according to the positive and negative controls as per the manufacturer’s instructions. The cells in the lower right quadrant were quantified as early apoptotic populations. C. Percentage of early apoptotic cells by PI-annexin V flow cytometry analysis during normal and HGPS iPSCs-keratinocytes induction. Data were presented as mean ± SD (n = 3). #P < 0.1, ns, not significant, Two-way ANOVA followed by Tukey’s multiple comparisons test.

Because the cell cycle abnormalities in differentiating HGPS epidermal cells resembles our previous findings in HGPS iPSCs derived smooth muscle cells (iSMCs), which is likely linked with mitotic catastrophe induced by progerin (Patel et al., 2021; H. Zhang et al., 2014), we next asked if there were increases in cell death during the HGPS iPSCs-keratinocytes induction. To address this question, we performed an Annexin V-PI assay using cells at the intermediate and late stages of differentiation. The population sorted as Annexin V-positive, and PI-negative (early apoptotic cells) was quantified as an indicator for cell death (Fig 3B). We observed that cell death in both normal and HGPS differentiation increased from the intermediate stage to the late stage. There was a slightly higher percentage of dying cells in HGPS iPSCs differentiation at each stage than normal control (Fig 3C).

### LEF1 is down-regulated in early HGPS iPSCs-keratinocytes differentiation

Previous studies identified WNT signaling as one of the most affected pathways in mouse skin keratinocytes with epidermal-specific progerin expression (Sola-Carvajal et al., 2019). WNT signaling is an early morphogenetic pathway that has been activated upon gastrulation and orchestrates multiple processes in the skin development (Fuchs, 2007; Lim & Nusse, 2013). Impaired canonical WNT pathway with diminished nuclear β-catenin localization has been reported in progeroid mouse skins and HGPS osteoblast differentiation (Choi et al., 2018; Sola-Carvajal et al., 2019). Given that, we would like to examine if the accelerated keratinocyte lineage-commitment in HGPS iPSCs differentiation comes along with an altered WNT pathway.

We first checked the β-catenin level by Western blot analysis in iPSCs-keratinocytes differentiation. We detected the weekly expression of the total and active form of β-catenin throughout the differentiation process (Fig 4A). Our results indicated that β-catenin expression was activated early upon the iPSCs-keratinocytes induction and was decreased overtime under normal and HGPS conditions. At each week, the β-catenin level in HGPS was quite comparable to normal conditions (Fig 4A and EV2A). Because progerin was reported to disrupt the nuclear translocation of β-catenin (Choi et al., 2018; Sola-Carvajal et al., 2019), we then performed nuclear extraction for the early-stage (week 1) differentiating cells with high β-catenin expression. Still, no significant difference in active β-catenin expression was detected in the HGPS nucleus (Fig 4B and EV2B). We then asked if the transcription of those essential WNT pathway genes were activated upon the early iPSCs-keratinocytes induction. Indeed, despite the unchanged TCF7 mRNA level, significant upregulations of LEF1, TCF7L1, and TCF7L2 transcription were detected at differentiation day 5, following the second stimuli with BMP-4 and RA. Interestingly, the LEF1 mRNA level in HGPS did not catch up with normal conditions at differentiation day 5, while no other gene expression showed the same trend (Fig 4C). We further detected the weekly expression of LEF1 throughout the differentiation process. Again, diminished LEF1 expression was observed in HGPS cells at all differentiation stages (Fig 4D and E).

**Figure 4:**
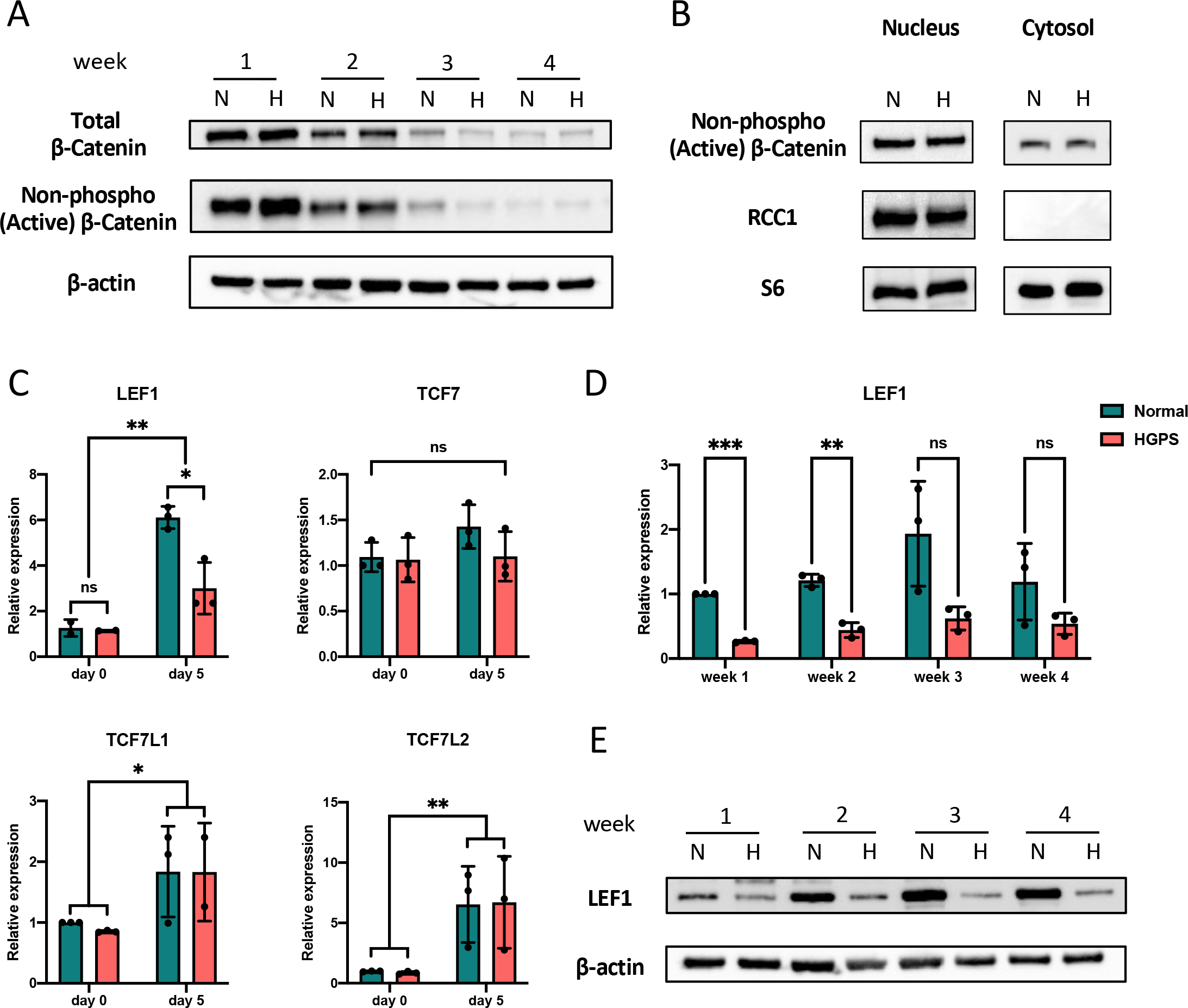
LEF1 down-regulation in early HGPS iPSCs-keratinocytes differentiation. A. Western blot analysis of total and non-phospho (active) β-catenin expression during normal and HGPS iPSCs-keratinocytes induction. Experiments were repeated at least three times, and representative data were shown as indicated. B. Western blot analysis of nuclear and cytosolic non-phospho (active) β-catenin level in early-differentiating (week 1) normal and HGPS iPSCs. RCC1 and S6 were used as a nuclear marker and the overall loading control, respectively. Experiments were repeated at least three times, and representative data were shown as indicated. C. Quantitative RT-PCR analysis of the relative expression of WNT transcription factors LEF1, TCF7, TCF7L1, and TCF7L2 before (day 0) and after induction with RA and BMP-4 (day 5) in normal and HGPS iPSCs differentiation. Data were normalized to endogenous *ACTB* mRNA and to the average of Normal day 0. Data were presented as mean ± SD (n = 3). \**P* < 0.05, \*\**P* < 0.01, ns, not significant, Two-way ANOVA followed by Tukey’s multiple comparisons test. D. Quantitative RT-PCR analysis of the relative expression of LEF1 during normal and HGPS iPSCs-keratinocytes induction. Data were normalized to endogenous *ACTB* mRNA and to the average of Normal week 1. Data were presented as mean ± SD (n = 3). \*\**P* < 0.01, \*\*\**P* < 0.001, ns, not significant, Two-way ANOVA followed by Sidak’s multiple comparisons test. E. Western blot analysis of LEF1 expression during normal and HGPS iPSCs-keratinocytes induction. Experiments were repeated at least three times, and representative data were shown as indicated.

### LEF1 regulates keratinocytes differentiation through K8/K18

To investigate if LEF1 is an upstream regulator that determines the iPSCs’ fate towards keratinocyte lineage upon RA and BMP-4 stimuli, we performed a genome-wide search for potential LEF1 binding targets among genes highly expressed in early epidermal development. At the gene locus of epidermal progenitor markers K8 and K18, adjacent to each other at chromosome 12, we identified seven putative LEF1 binding sites (Fig 5A). We then performed a Chromatin immunoprecipitation (ChIP) assay using LEF1 antibody and quantified the percent input of these binding sites by qPCR. Our results indicated strong LEF1 recruitment at binding sites K8-1, K8-5, and K8-6, which correlates with LEF1 ChIP-seq peaks in HEK293T and human embryonic stem cells (HUES64) derived mesoderm (Fig 5B and EV3A). LEF1 binding at K8-1 was the strongest among them compared to a positive control at the *MYC* gene (Fig 5B).

**Figure 5:**
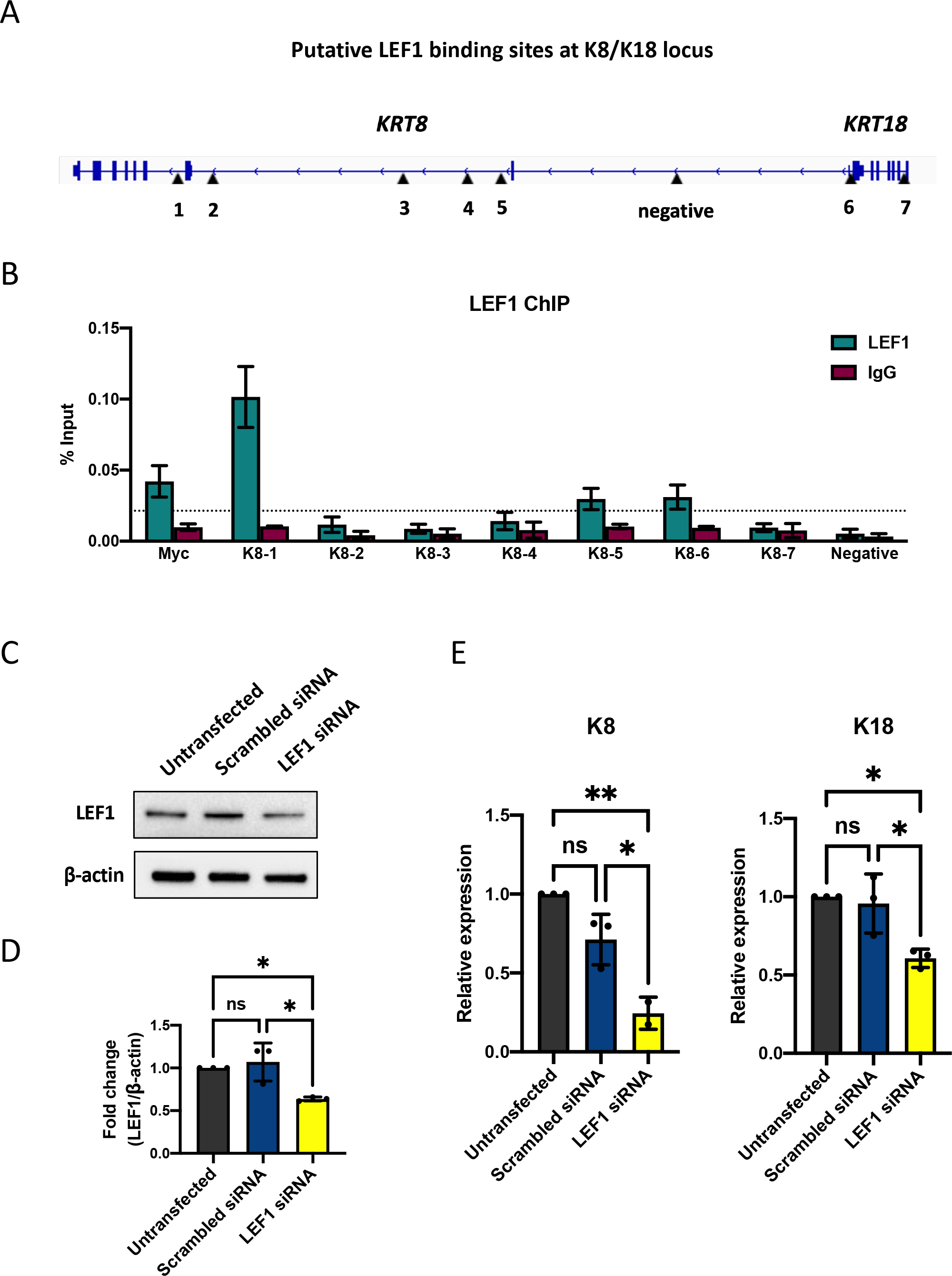
LEF1 regulates keratinocytes differentiation through K8/K18. A. Schematic representation showing putative LEF1 binding sites at K8/K18 gene locus. The position of seven putative LEF1 binding sites and a non-specific locus (negative) were indicated by black arrowheads. B. Chromatin immunoprecipitation quantitative PCR (ChIP-qPCR) analysis of the DNA binding activity of LEF1 at K8/K18 locus. A known LEF1 binding site at the MYC gene was used as a positive control. The binding sites with LEF1 enrichment 4 fold greater than the non-specific locus (above the dashed line) were considered to have strong LEF1 binding. Data were presented as mean ± SD (n = 3). C. Representative Western blot result showing LEF1 protein expression 48 hours after LEF1 siRNA knockdown in normal iPSCs differentiation. D. Quantification of LEF1 siRNA knockdown Western blot analysis in (C). Data were presented as mean ± SD (n = 3). \**P* < 0.05, ns, not significant, One-way ANOVA followed by Tukey’s multiple comparisons test. E. Quantitative RT-PCR analysis of the relative expression of K8 and K18 48 hours after LEF1 siRNA knockdown in normal iPSCs differentiation. Data were normalized to endogenous *ACTB* mRNA and to the average of untransfected samples. Data were presented as mean ± SD (n = 3). \**P* < 0.05, \*\**P* < 0.01, ns, not significant, One-way ANOVA followed by Tukey’s multiple comparisons test.

To further test the impact of LEF1 on K8/K18 expression, we down-regulated LEF1 expression at differentiation day 5 using a combination of three LEF1-targeting siRNAs. Our results showed that with a partial knockdown of LEF1 (Fig 5C and D), both K8 and K18 mRNA transcription decreased significantly (Fig 5E). Changes in their protein expression were not detected at this time point based on the Western blot analysis (Figure EV3B and C).

### Correcting the HGPS mutation in iPSCs-keratinocytes differentiation with ABE

To revert the *LMNA* c.1824 C>T mutation and curb its impact on HGPS iPSCs-derived keratinocytes from an early developmental stage, we conducted Adenine base editing (ABE) using the c.1824 C>T targeting ABE7.10max lentiviral vector at differentiation day 5, followed by three days of puromycin selection. ABE7.10max vector with human non-targeting sgRNA was included (mock-corrected) (Fig 6A). At the end of iPSCs-keratinocytes differentiation (week 4), about 50% of *LMNA* c.1824 T (pathogenic) was corrected to C (wild-type) (Fig 6B). Although progerin was still detectable in HGPS-corrected iPSCs-keratinocytes, immuno-stained cells with very bright nuclei consisting of highly expressed progerin were rarely seen after ABE correction (Fig 6C). Our RT-qPCR results showed a 50% reduction of progerin mRNA transcription in HGPS- corrected iPSCs-derived keratinocytes (Fig 6D) and that the ABE treatment led to a significant decrease in progerin expression (Fig 6E). Interestingly, the keratinocyte markers expression in HGPS-corrected cells was significantly reversed despite the moderate correction efficiency.

**Figure 6.**
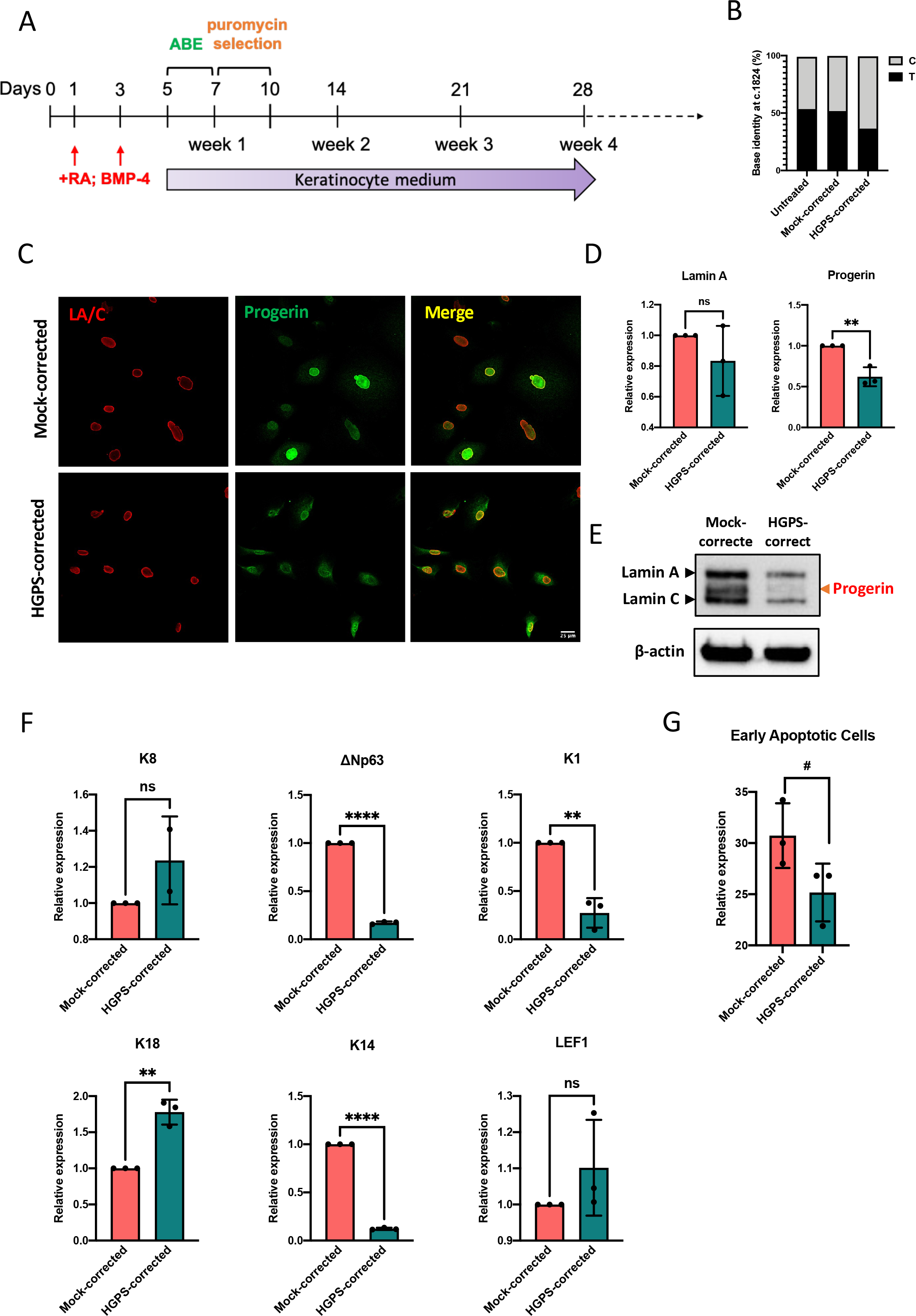
ABE corrects the HGPS mutation in iPSCs derived keratinocytes. A. Schematic representation of the Adenine base editing in HGPS patient-derived iPSCs-keratinocytes induction. B. *LMNA* c.1824 nucleotide identity in HGPS iPSCs derived-keratinocytes untreated or treated with ABE7.10max-VRQR lentivirus (mock and HGPS mutation-targeting) at differentiation week 4. C. Lamin A/C and progerin expression in mock-corrected and HGPS-corrected iPSCs-derived keratinocytes at week 4 indicated by immunofluorescence staining (scale bar = 25μm). D. Quantitative RT-PCR analysis of the relative expression of Lamin A and progerin in mock-corrected and HGPS-corrected iPSCs-derived keratinocytes at week 4. Data were normalized to endogenous *ACTB* mRNA and to the average of mock-corrected cells. Data were presented as mean ± SD (n = 3). \*\**P* < 0.01, ns, not significant, unpaired two-tailed *t*-test. E. Western blot analysis of Lamin A/C and progerin expression in mock-corrected and HGPS-corrected iPSCs-derived keratinocytes at week 4. Experiments were repeated at least three times, and representative data were shown as indicated. F. Quantitative RT-PCR analysis of the relative expression of ΔNp63, K14, K8, K18, K1, and LEF1 in mock-corrected and HGPS-corrected iPSCs-derived keratinocytes at week 4. Data were normalized to endogenous *ACTB* mRNA and to the average of mock-corrected cells. Data were presented as mean ± SD (n = 3). \*\**P* < 0.01, \*\*\*\**P* < 0.0001, ns, not significant, unpaired two-tailed *t*-test. G. Percentage of early apoptotic cells by PI-annexin V flow cytometry analysis in mock-corrected and HGPS-corrected iPSCs-derived keratinocytes at week 4. Data were presented as mean ± SD (n = 3). #*P* < 0.1, unpaired two-tailed *t*-test.

Down-regulated p63, K14 and K1 and up-regulated K8 and K18 expression were detected after the base editing (Fig 6F and EV4A). At differentiation week 4, the mRNA transcription of LEF1 was slightly restored in HGPS-corrected iPSCs-keratinocytes (Fig 6F). Moreover, the phenotype of massive cell death was alleviated in HGPS-corrected iPSCs-derived keratinocytes (Fig 6G), although the cell cycle partitioning remained unchanged (Fig EV4B).

## Discussion

### Stem cell depletion may contribute to the accelerated keratinocyte-commitment in HGPS iPSCs differentiation

In 2007, Halascheck-Wiener *et al*. promoted a stem cell exhaustion theory to explain the segmental aging phenotypes that appears to be more severe in tissues under continuous strong mechanical stress or endure high turnover rate in HGPS (Halaschek-Wiener & Brooks-Wilson, 2007). For instance, the nuclear accumulation of progerin in HGPS endothelial cells and smooth muscle cells induces genome instability and increases cell death and turnover rate, resulting in a depletion of endothelial progenitor cells (EPCs). The continuing loss of EPCs will impair vascular maintenance and regeneration, causing severe cardiovascular conditions (Caplice & Doyle, 2005; Roberts, Jahangiri, & Xu, 2005). The stem cell exhaustion theory may also explain the hair loss and lipodystrophy in HGPS (Halaschek-Wiener & Brooks-Wilson, 2007). Moreover, epidermal hyperplasia and premature depletion of adult stem cells were reported in mouse skins with tissue-specific progerin expression (Rosengardten et al., 2011). In this study, we observed an accelerated commitment of HGPS iPSCs to keratinocyte lineage, based on the keratinocyte development markers at early, middle and late stages. Consistent with studies in other HGPS cell types, these differentiating HGPS keratinocytes showed a higher percentage of G2/M cells and an increased apoptosis rate. Together, these results suggest that HGPS keratinocytes differentiation is accelerated, resulting in undesired, hastened cell turnover and eventual depletion of epithelial stem cells.

### LEF1 is an early regulator in epidermal development, and its down-regulation accelerates the keratinocyte lineage commitment in HGPS

WNT signaling has been identified as one of the most affected pathways in the HGPS mouse keratinocytes (Sola-Carvajal et al., 2019). In iPSCs-keratinocytes differentiation, many WNT pathway components were highly expressed upon the induction of RA and BMP-4. Interestingly, we found that the expression of a critical WNT transcription factor LEF1 was diminished at an early stage in HGPS iPSCs differentiation. Moreover, the HGPS iPSCs-derived cells maintained a lower LEF1 expression level throughout the epidermal development than the normal control. We further investigated the impact of LEF1 on the expression of epidermal progenitor genes K8 and K18. In iPSCs-keratinocytes differentiation, K8/K18 expression was induced upon RA treatment and decreased over time significantly after p63 was activated (L. Zhang, 2018). In HGPS iPSCs-keratinocytes induction, the expression of K8/K18 decreases more rapidly, possibly due to insufficient LEF1 level, thus resulting in an accelerated commitment towards the p63 and K5/K14-positive keratinocyte lineage (Fig 7B).

**Figure 7.**
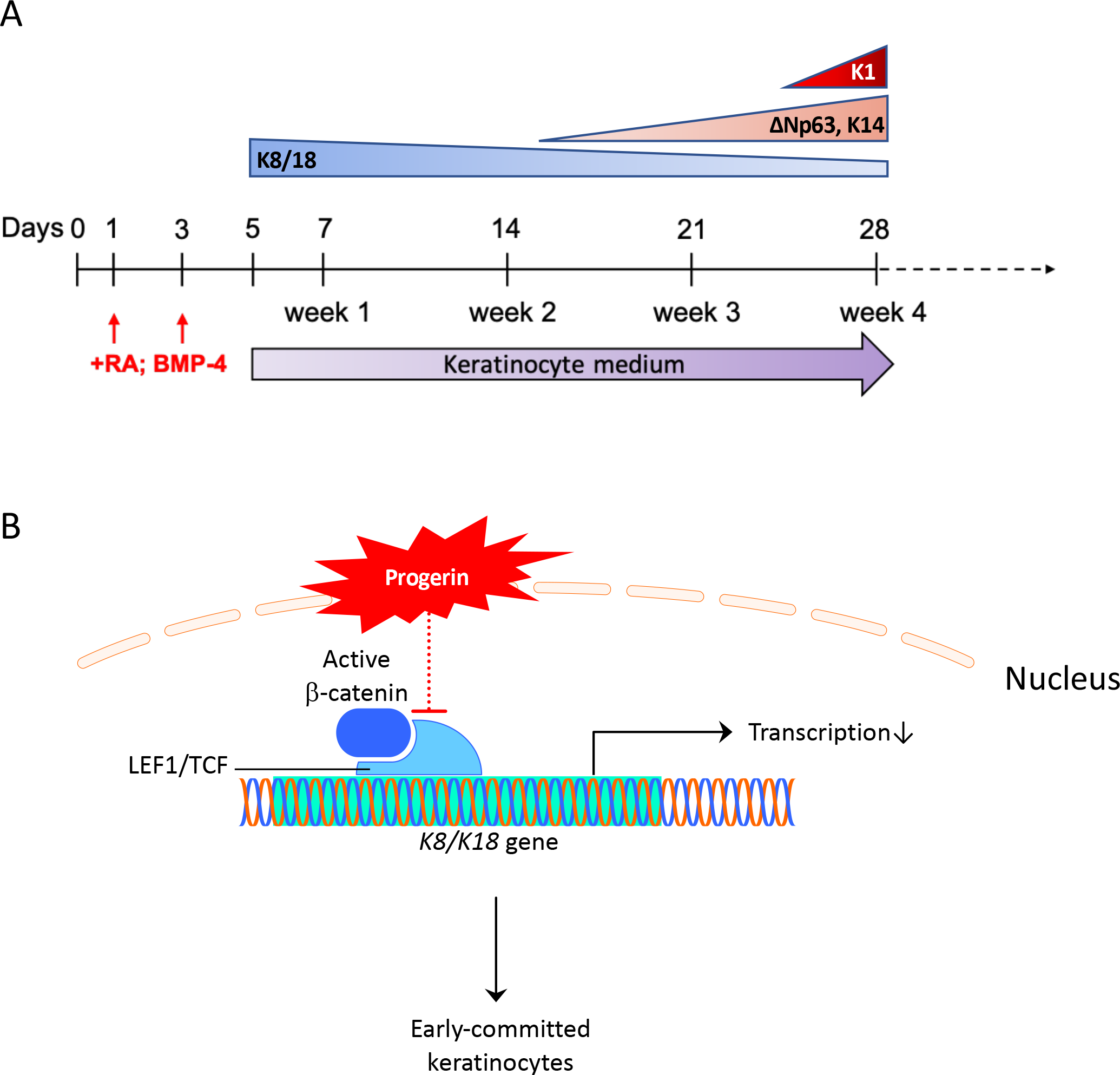
Working model of epidermal development in HGPS. A. Schematic representation of the keratinocyte-related markers succession during iPSCs- keratinocytes induction. B. Schematic diagram showing the molecular mechanism for the early commitment to the keratinocyte lineage in HGPS iPSCs differentiation.

Previous studies revealed a β-catenin loss in the HGPS cell nucleus due to disrupted nuclear transportation caused by the progerin (Choi et al., 2018; Sola-Carvajal et al., 2019). In our iPSCs- keratinocytes differentiation process, the nuclear β-catenin level was unchanged at the early stage of iPSCs-keratinocytes differentiation, when progerin aggregates were not yet formed at the nuclear envelope. Although it is unclear how LEF1 transcription was inhibited at this stage of differentiation, other potential pathways regulate LEF1 expression in a β-catenin-independent manner, including the BMP pathway, which activates LEF1 transcription via P-SMAD in the neural stem cells differentiation (Armenteros, Andreu, Hortigüela, Lie, & Mira, 2018).

### ABE in HGPS iPSCs-keratinocytes differentiation

Since the developmental defects in HGPS iPSCs-keratinocytes have an early onset, we strived to correct the c.1824C>T mutation at an early developmental stage without disturbing the initial inductions with RA and BMP-4. Our adenine base editing in HGPS iPSCs-derived keratinocytes did not reach the high correction efficiency of primary fibroblasts or endothelial cells. As a result, approximately 30% of the cells still carry the HGPS mutation at differentiation day 28. Notably, in HGPS animals with embryonic ABE-AAV9 virus injection, the base editing efficiency in the skin was also lower than in other tissues such as the liver and heart (Koblan et al., 2021). One possible explanation lies in the cell cycle abnormalities in HGPS epidermal development. Because progerin was relatively stable, it may persist in the HGPS-corrected cells long after the mutation has been corrected. Thus, it may take a long period for complete progerin clearance.

However, as basal layer keratinocytes have limited self-renewal potential and start spontaneous terminal differentiation under the *in vitro* culture condition, there was only a short window to evaluate the properties of HGPS-corrected cells. Based on our cell cycle analysis, progerin left a constant impact on the HGPS-corrected cell division. According to the cell cycle partitioning in normal and HGPS cells, at differentiation week 2, when ABE had its optimal effects, reducing the S-phase population made it difficult for genome editing to occur. Another unique feature in epidermal development may also reduce the base editing efficiency: the high cell death and turn-over rate in normal and HGPS differentiation. ABE only slightly reduced the cell death in HGPS-corrected cells. Interestingly, our results indicated that even a tiny portion of correction makes a big difference in decelerating the commitment towards keratinocytes in HGPS. A similar phenomenon was seen in ABE-corrected HGPS animals.

Despite the small percentage of correction in many tissues, their health condition improved greatly with significantly extended lifespan (Koblan et al., 2021). At the end of differentiation, only a slightly higher LEF1 expression was detected, indicating a less critical role of LEF1 at the late stage of epidermal development. Future experiments such as ABE in HGPS iPSCs may increase the correction efficiency in HGPS iPSCs-derived keratinocytes.

## Materials and Methods

### Keratinocytes differentiation from iPSCs

Normal and HGPS patient-derived iPSCs used in this study were reprogramed from skin fibroblasts of a male HGPS patient who carried the classic G608G HGPS mutation (HGADFN167) and his normal father (HGADFN168) and provided by Progeria Research Foundation Cell and Tissue Bank. *In vitro* differentiation protocol towards the keratinocyte lineage was adapted from previous publications with minor modifications (Itoh et al., 2011; Kogut et al., 2014). In brief, feeder-free iPSCs were cultured in defined keratinocytes serum-free medium (DKSFM, Gibco) containing 25 ng/ml bone morphogenetic protein-4 (BMP-4, R&D Systems) and 1 μM all-trans Retinoid Acid (RA, Sigma-Aldrich) at the initial stage of differentiation (medium was changed once at differentiation day 3). Starting from day 5, DKSFM was changed every other day for up to 28 days. Fully differentiated keratinocytes can be passed 3-5 times on Collagen I (Sigma-Aldrich)-coated tissue culture dishes. Cells were maintained at 37℃ and 5% CO^2^ in a humidified incubator.

### Adenine base editor lentiviral vectors production

The ABE7.10max-VRQR lentiviral vector targeting the HGPS G608G mutation was generated as described in Koblan *et al*., 2021. Human non-targeting control vectors were generated as described in Gete *et al*., 2021.

### Immunocytochemistry

For immunocytochemistry, cells were washed twice with PBS and fixed in 4% paraformaldehyde (PFA) for 15 mininutes, followed by permeabilization with 0.5% Triton in PBS for 5 minutes. Cells were then blocked with 4% BSA in PBS for one hour. After that, cells were incubated with primary antibodies in 4% BSA in PBS overnight at 4°C. Next day, after five PBS washes, cells were incubated in secondary antibodies diluted in 4% BSA in PBS for one hour. Cells were then washed five times with PBS, and fluorescence images were acquired with Zeiss LSM 710 confocal microscope (Zeiss International, Oberkochen, Germany). Fluorescence intensity was adjusted with ImageJ software (NIH). Primary antibodies used for immunocytochemistry were: lamin A/C (1:250, Millipore MAB3211), progerin (1:250, Cao et al., 2011), ΔNp63 (1:250, Biolegend #619001), K14 (1:500, Invitrogen MA5-11599). Secondary antibodies used were: Alexa Fluor 488 donkey anti-rabbit IgG (1:1000, Invitrogen), Alexa Fluor 594 donkey anti-rabbit IgG (1:1000, Invitrogen), Alexa Fluor 488 donkey anti-mouse IgG (1:1000, Invitrogen), and Alexa Fluor 594 donkey anti-mouse IgG (1:1000, Invitrogen).

### Western blot

Whole-cell lysates for immunoblotting were prepared by dissolving cells in Laemmli Sample Buffer containing 5% 2-mercaptoethanol (Bio-Rad). Nuclear and cytosolic fractions of iPSCs-differentiated cells were acquired using NE-PER™ Nuclear and Cytoplasmic Extraction Reagents (Thermo Scientific #78833) as per the manufacturer’s instructions. Protein samples were loaded on 10%-12% polyacrylamide gels and transferred onto 0.45 µm pore-size nitrocellulose membranes (Bio-Rad) using the Turboblot (BioRad). Blots were incubated overnight at 4°C with primary antibodies and then probed with secondary antibodies before ECL development and imaging (Bio-Rad). Primary antibodies used for immunoblotting are as follows: lamin A/C (1:500, Millipore MAB3211), progerin (1:500, Cao et al., 2011), ΔNp63 (1:500, Biolegend #619001), K14 (1:500, Invitrogen MA5-11599), K8 (1:500, Biolegend #904804), K18 (1:1000, Cell Signaling #4548), total β-Catenin (1:1000, Cell Signaling #9562), Non-phospho (Active) β-Catenin (1:1000, Cell Signaling #8814), RCC1 (1:1000, Cell Signaling #3589), S6 Ribosomal Protein (1:1000, Cell Signaling #2217), LEF1 (1:250, Santa Cruz sc-374412) and β-actin (1:1000, Sigma-Aldrich A3854).

### RNA isolation and Real-Time quantitative PCR

Total genomic RNA was extracted with Trizol (Life Technologies) and purified using the RNeasy Mini kit (Qiagen) as per the manufacturer’s instructions. The RNA yield was determined by the NanoDrop 2000 spectrophotometer (Thermo Fisher). 1 µg of total RNA was converted to cDNA using iScript Select cDNA Synthesis kit (Bio-Rad). Quantitative RT-PCR was performed in triplicate using SYBR Green Supermix (Bio-Rad) on CFX96 Real-Time PCR Detection System (C1000 Thermal Cycler, Bio-Rad). All primers used in this study are listed in Table EV1.

### Cell cycle analysis

Cell cycle analysis was performed at differentiation week 2 and week 4 in normal and HGPS iPSCs-keratinocytes induction. Before harvest, cells were incubated in 10 uM BrdU (BD #550891) labeling medium for one hour. The collected cells were then washed once and resuspended in 100 μL PBS. The cell suspension was fixed in 1 mL of 70% (vol/vol) ethanol for one hour on ice, followed by a PBS wash and denatured in 2N HCl with 0.5% Triton X-100 for 30 minutes at RT. Cells were washed again with PBS, blocked and incubated in Alexa Fluor 488 anti-BrdU antibody solution (1:100, Invitrogen #B35130) at 4 °C overnight. On the next day, the cells were pelleted, rinsed with PBS, and resuspended in 50 μg/mL propidium iodide (PI, Invitrogen) with 100 μg/mL DNase-free RNase (Thermo Scientific) in PBS for 30 minutes at 37 °C. Flow cytometry was performed with FACS CantoII (BD) and the data were analyzed by FlowJo software.

### Apoptosis assay

PI-annexin V apoptosis assay was performed at differentiation week 2 and week 4 in normal and HGPS iPSCs-keratinocytes induction according to the manufacturer’s instruction (BD). In brief, cells were harvested and rinsed with PBS, then resuspended and stained with 100 μL of 1x annexin V binding buffer containing 5 μL of annexin V and 5 μL of PI for 25 minutes in the dark at room temperature. Stained samples were analyzed by FACS CantoII (BD) and the data were processed by FlowJo software.

### Chromatin immunoprecipitation

LEF1 ChIP was performed in early-differentiating (week 1) epidermal cells derived from normal iPSCs. Detailed experimental procedures were described in the protocol of SimpleChIP® Plus Sonication Chromatin IP Kit (Cell Signaling #56383). Briefly, protein and DNA were cross-linked using 1% formaldehyde for 10 minutes. Nuclei were isolated and lysed, and chromatin was sheared to an average size of 400 bp by sonication. A small aliquot of the supernatant was used as input control, and the remaining sonicated chromatin was divided into two aliquots, which were incubated with an anti-LEF1 antibody (1:50, Cell Signaling #76010) and anti-rabbit IgG antibody (1:1000, or 1 μL for one immunoprecipitation, Cell Signaling #2729) as a negative control, respectively. The antibody-bound complex was precipitated with Dynabeads protein G (Invitrogen #10004D). The DNA fragments were released from the immunoprecipitated complexes by reversing the cross-linking at 65°C overnight. The precipitated DNA was isolated, purified and used as a template for PCR. Primers used for ChIP-qPCR were listed in Table EV1.

### Silencing experiments

The siRNA plasmids consist of a pool of three human LEF1-targeting sequences (Santa Cruz, sc-35804) and the control scrambled siRNA (Santa Cruz, sc-37007) were purchased from Santa Cruz. Transient transfections were carried out using Lipofectamine RNAiMAX Transfection Reagent (Invitrogen) following the manufacturer’s protocol. After 48 h of transfection, cells were collected for RT-qPCR and Western blotting analysis.

### Statistical analysis

Statistical analyses were performed using GraphPad Prism 7 software. Data were analyzed using unpaired Student’s *t*-test for two groups. One-way and two-way analysis of variance (ANOVA) followed by post hoc multiple comparisons were used to compare means of three or more groups. Analysis method adapted in each experiment was indicated in figure legends. All experiments were repeated at least three times, and results were presented as the mean ± SD. *p* value < 0.05 or < 0.1 was considered significant.

### Data availability

This study includes no data deposited in external repositories.

## Acknowledgments

We thank Drs. John Fisher, Margaret Scull, Sougata Roy, and Jiqiang Ling from the University of Maryland for providing insightful feedback to this work. We thank the Imaging Core and FACS core at University of Maryland for their technical support. We would like to thank all the members from Cao lab for their helpful discussions and suggestions.

## Author contributions

XM and KC designed the research; XM, ZMX, HX, MAB, YGG, and LS conducted the research; XM and RY analyzed the data; XM collected the data, generated, and compiled figures, and wrote the manuscript; KC reviewed the manuscript. All authors read and approved the final manuscript.

## Supporting Information

**Figure EV1.**
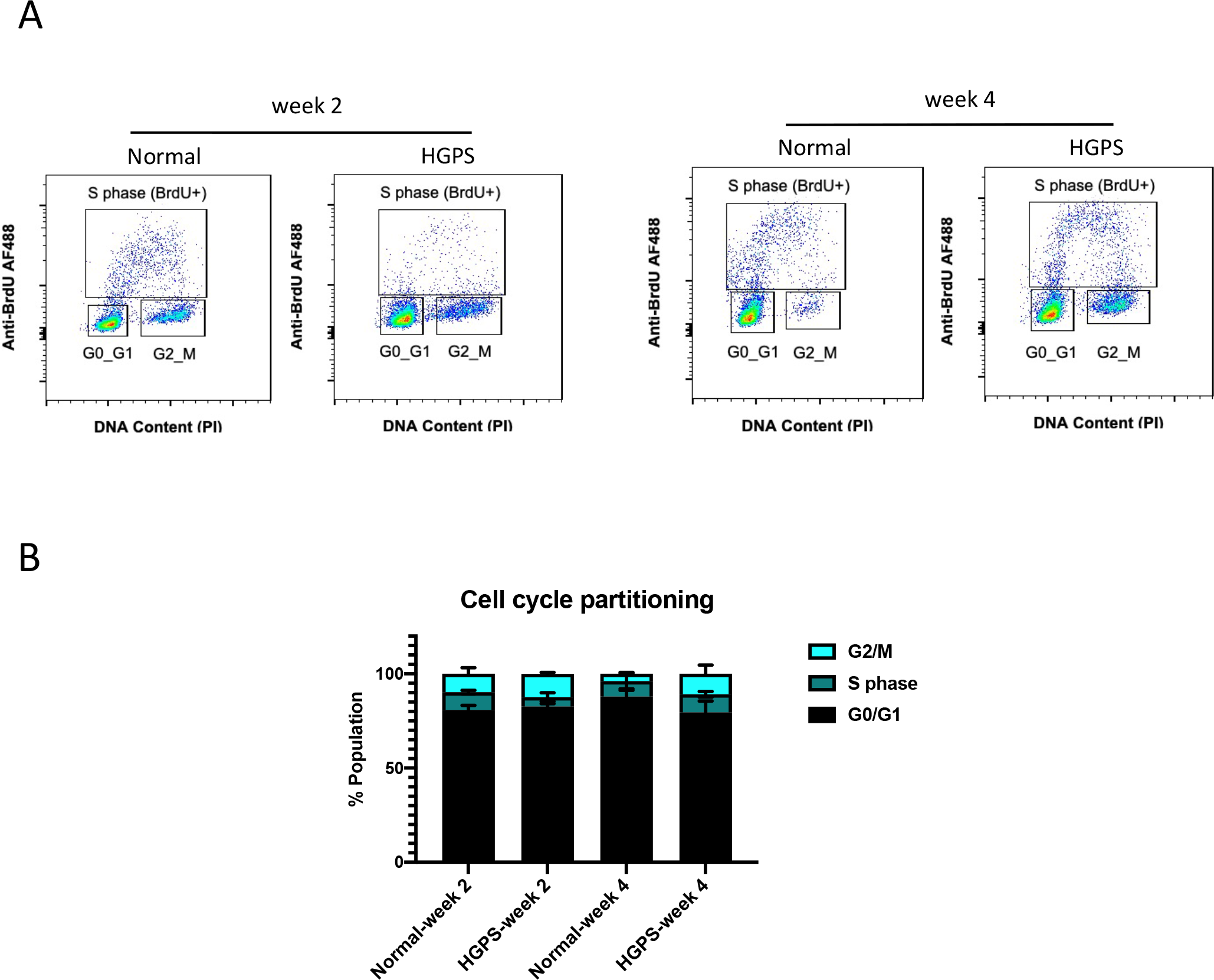
A. Representative flow cytometry plots showing BrdU-PI cell cycle analysis during normal and HGPS iPSCs-keratinocytes induction. The distribution of cells at G0/G1, S, and G2/M phases were indicated within each plot. B. Quantification of the cell cycle partitioning during normal and HGPS iPSCs-keratinocytes induction. Data were presented as mean ± SD (n = 3).

**Figure EV2.**
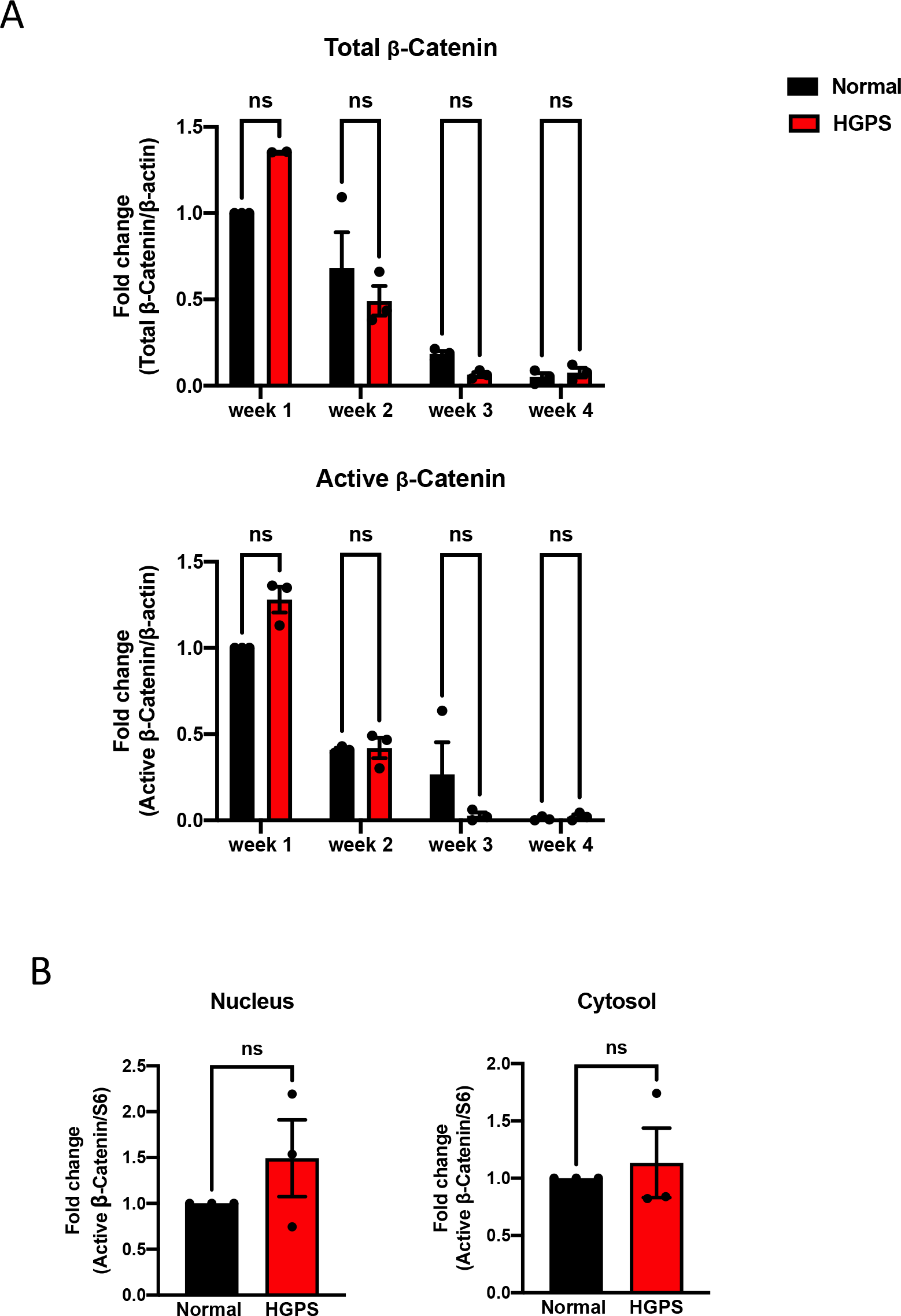
A. Quantification of total and active β-catenin expression during normal and HGPS iPSCs-keratinocytes induction. The Representative Western blot result is shown in Figure 4A. Data were presented as mean ± SD (n = 3). ns, not significant, Two-way ANOVA followed by Sidak’s multiple comparisons test. B. Quantification of nuclear and cytosolic active β-catenin expression in early-differentiating (week 1) normal and HGPS iPSCs. The Representative Western blot result is shown in Figure 4B. Data were presented as mean ± SD (n = 3). ns, not significant, unpaired two-tailed *t*-test.

**Figure EV3.**
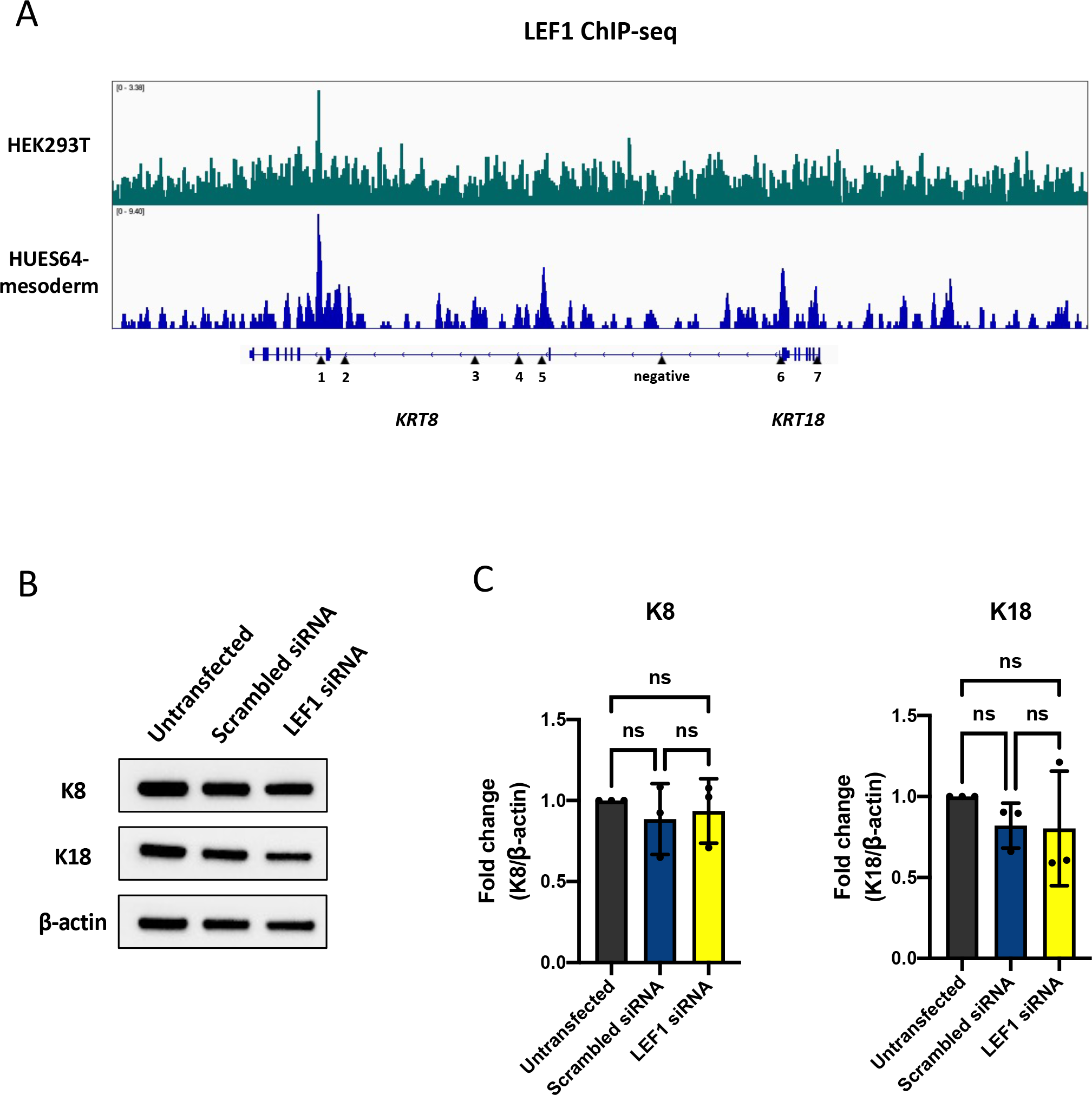
A. Genomic tracks display public available LEF1 ChIP-seq data. Shown are LEF1 enrichment profiles at K8/K18 locus in HEK293T (green) and human embryonic stem cell (HUES64) derived mesoderm (blue). The K8/K18 gene track with putative LEF1 binding sites was shown below the profiles. B. Representative Western blot result showing K8 and K18 protein expression 48 hours after LEF1 siRNA knockdown in normal iPSCs differentiation. C. Quantification of K8 and K18 protein expression after LEF1 siRNA knockdown in (B). Data were presented as mean ± SD (n = 3). ns, not significant, One-way ANOVA followed by Tukey’s multiple comparisons test.

**Figure EV4.**
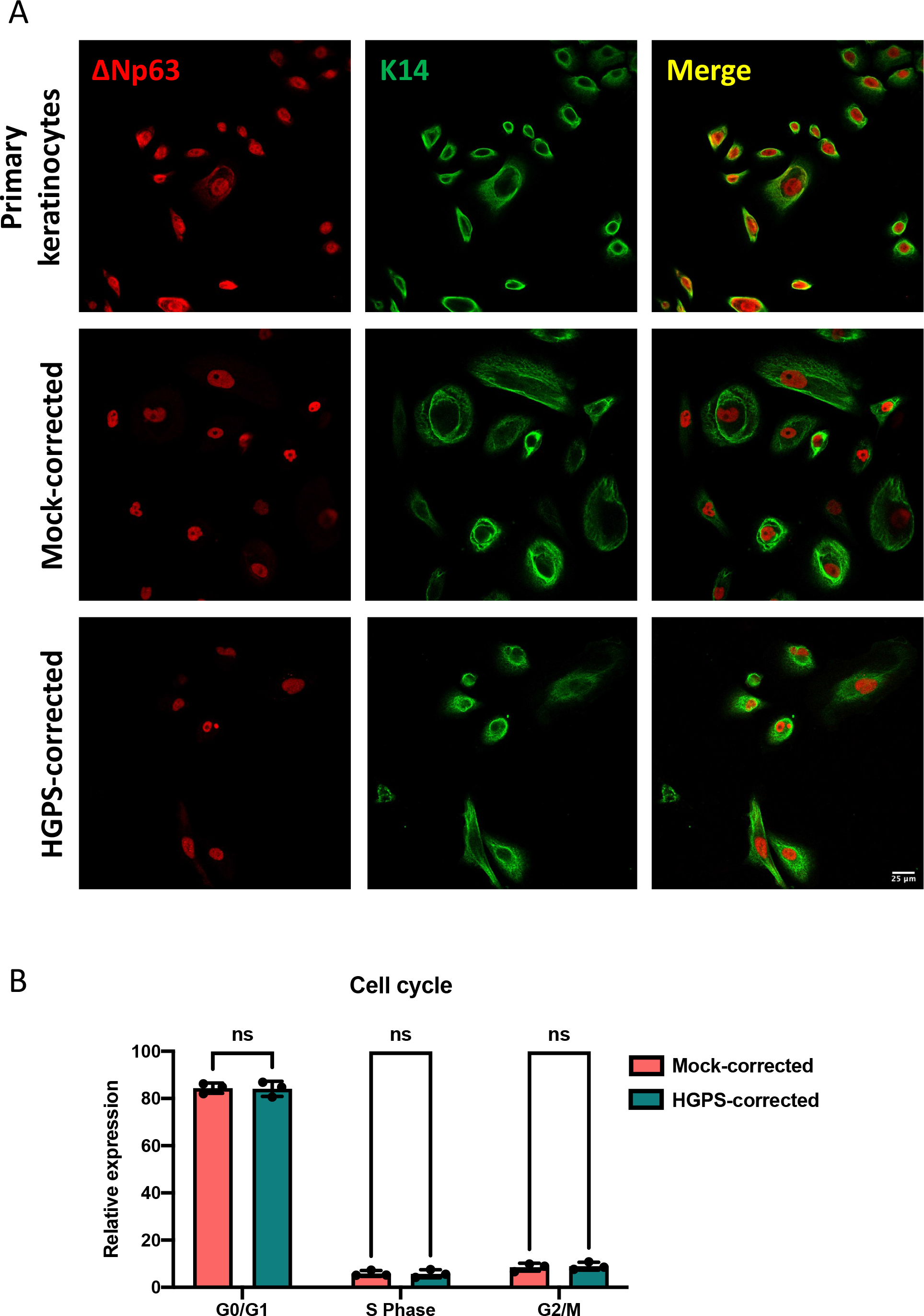
A. Basal layer keratinocyte markers ΔNp63 and K14 expression in mock-corrected and HGPS-corrected iPSCs-derived keratinocytes at week 4 as well as in primary keratinocytes indicated by immunofluorescence staining (scale bar = 25μm). B. Quantification of the cell cycle partitioning in mock-corrected and HGPS-corrected iPSCs-derived keratinocytes at week 4. Data were presented as mean ± SD (n = 3). ns, not significant, Two-way ANOVA followed by Sidak’s multiple

